# VGLUT2/ Cdk5/p25 Signaling Pathway Contributed to Inflammatory Pain By CFA

**DOI:** 10.1101/655498

**Authors:** YuWen Tang, ZhiYou Peng, ShouJun Tao, Jianliang Sun, WenYuan Wang, XueJiao Guo, GongLu Liu, XianZhe Luo, Yuan Chen, Yue Shen, HaiXiang Ma, Peng Xu, Qing Hua Li, HongHai Zhang, ZhiYing Feng

## Abstract

Vesicular glutamate transporter type 2 (VGLUT2) is known to play an important role in mediating the heat hyperalgesia induced by inflammation. However, the underlying mechanism for this activity is poorly understood. Cyclin-dependent kinase 5 (Cdk5), serving as a key regulator in mediating release of glutamate, contributed to the inflammatory heat. It remains unknown whether there is a bridge between Cdk5 and VGLUT2 for mediating inflammatory pain. Therefore, we designed the experiment to determine whether VGLUT2 signaling pathway is involved in Inflammatory pain mediated by Cdk5 and the heat hyperalgesia induced by complete Freund’s adjuvant (CFA) can be reversed by roscovitine, a selective inhibitor for Cdk5 through inhibition of VGLUT2 expression. Immunohistochemistry results suggest that when compared with rats in a control group, rats in an experimental group showed significant coexpression of Cdk5/VGLUT2 in small and medium-sized neuronal cells of the dorsal root ganglion (DRG) and spinal cord between days 1 and 3 following subcutaneous injection of CFA. Moreover, our study revealed that the expression of VGLUT2 protein in DRG and spinal cord was remarkably increased between days 1 and 3 following CFA injection. Additionally, p25 but not p35, a activator of Cdk5, protein was significantly increased and reduced by roscovitine. The increased expressions of VGLUT2 protein was significantly reduced by roscovitine as well. Our study showed that VGLUT2/Cdk5 signaling pathway contributed to the inflammatory pain medicated by Cdk5/p25.

## Introduction

Pain caused by inflammation in the peripheral or central nervous system remains a significant clinical problem, and is often resistant to treatment with conventional analgesics. Glutamate is the major excitatory neurotransmitter in the central nervous system (CNS) and plays a key role in processing of nociceptive pain [1]. Prior to its release from excitatory synapses, glutamate is transported into synaptic vessels by vesicular glutamate transporters (VGLUTs) composed of VGULT proteins 1-3. Previous studies have established that VGLUT1, 2, and 3 proteins are expressed by largely non-overlapping and functionally distinct populations of glutamatergic neurons, and play multiple roles in the CNS [2-4]. However, only VGLUT 2 plays a key role in mediating pain hypersensitivity induced by inflammation and peripheral nerve injury [5-12]. Co-expression of VGLUT2 and two main nociceptors of calcitonin gene related peptide (CGRP) and transient receptor potential channel (TRPV1) was observed in small and medium-sized neurons of the dorsal root ganglion (DRG) in mice [6, 7]. Additionally, VGLUT2 knock-out mice challenged with CFA demonstrate a complete loss of heat hyperalgesia, whereas CFA-induced mechanical hypersensitivity remains intact. Nevertheless, a detailed mechanism for VGLUT2 mediation of inflammation–induced heat hyperalgesia remains elusive.

Emerging evidence suggests that the serine/threonine kinase Cdk5 plays an important role in mediating inflammation-induced heat hyperalgesia [13-16]. A previous study showed that Cdk5 can modulate CFA-induced heat hyperalgesia by controlling membrane trafficking of TRPV1 and phosphorylating vanilloid receptor 1 (VR1) in the dorsal root ganglion (DRG) of rats [14, 15]. The activities of Cdk5 and p35 in the spinal cord were significantly increased following peripheral injection of CFA. Furthermore, both Cdk5 kinase activity and heat hyperalgesia were inhibited by Cdk5/p35 knockdown or intrathecal administration of roscovitine. Previous studies have suggested that presynaptic Cdk5 is the primary regulator of neurotransmitter release in the CNS [17]. Synapsin II is a synaptic vesicle protein associated with the release of neurotransmitters, and a recent study revealed that expression of both VGLUT1 and VGLUT2 was strongly reduced in the spinal cord of synapsin II knockout mice [18]. Moreover, our previous studies revealed that increased levels of synaptophysin protein, an important presynaptic vesicle membrane protein that functions in release of neurotransmitters, are involved in mediating CFA-induced heat hyperalgesia mediated by Cdk5 in rats. Furthermore, the recent study showed that VGLUT2-pH fluorescence co-localizes with synaptophysin at synaptic boutons and was invovled in the trafficking of synaptic vesicles [19]. This suggests that Cdk5 may mediate inflammation-induced heat hyperalgesia by controlling the release of neurotransmitters [20]. Here, we report that the VGLUT2/Cdk5 signaling pathway contributed to the inflammatory pain medicated by Cdk5. Materials and methods

### Animals

All adult male Sprague-Dawley rats (200-250g) used in this study were obtained from the animal center of Nanjng Medical University (Nanjing China). All experimental procedures were verified and approved by the Committee of Animal Use for Research and Education of Nanjing Medical University. Moreover, these procedures were performed in accordance with guidelines developed by the International Association for the Research on Pain [21]. According to the guidelines mentioned before, rats were placed with room temperature of 22 ± 2°C and a standard 12/12 hours light/dark cycle, and food and water were available ad libitum. To minimize the animals we used in this research and avoid any unnecessary stress and pain, all animals were permitted to adapt to the housing facilities for 1 week before the experiments. Complete Freund’s Adjuvant CFA (100 µL; Sigma, St. Louis, MO, USA) and saline (100 µL) were injected into the plantar surface of the ipsilateral and contralateral hind paws of rats respectively. The saline group was settled as a control (n = 6/group). The same volume of saline as used for control group was injected into ipsilateral and contralateral paws of rats in the control group (n = 6/group).

### Surgery and drug administration

Drugs used in this study were intrathecal injection as described before, with a slight modification [22]. Briefly, a catheter (PE-10: 0.28 mm i.d. and 0.61 mm o.d.; Clay Adams, Parsippany, NJ, USA) was inserted at the lumbar level of the spinal cord between lumbar vertebrates 4 and 5 (L4 and L5) of the rats which were anesthetized before with the 4% pentobarbital (40 mg/kg). 5 µL of 2% lidocaine was confirmed to be injected through the catheter for the recovery after the anesthesia and surgery, followed by flushing with 15 µL of saline. Following this procedure, successful insertion of the catheter was further confirmed if the animal presented with impaired motor function of their hind legs at 10 s post-lidocaine administrations. Following surgery, 5 µL of roscovitine (100 µg) (Sigma, St. Louis, MO, USA R7772) dissolved in 10% DMSO was delivered for 5 consecutive days to the rats in the experimental group, followed by rinsing of 15 µL of sterile saline 0.5 h before a single treatment with CFA (n = 6/group). The same concentration of roscovitine and DMSO was injected into the rats in the vehicle control group (n = 6) and the experimental group.

### Behavioral test

Heat hyperalgesia was quantified by measuring paw withdrawal latencies (PWL) in response to radiant heat stimulation as previously described [23]. Briefly, radiant heat was directed to the plantar surface of each hind paw through a 1 mm thick glass plate. In order to prevent tissue damage, a 20 second cut-off limit was radicate for heat exposure. The time between the onset of the heat stimulus and a manifestation of paw withdrawal response was recorded as the thermal nociceptive latency period. Rats were allowed to adapt to the room for 30 min before the behavioral test. The thermal stimulus was then delivered to the ipsilateral and contralateral hind-paws of rats in both groups after adaption. The time of PWL measurements was at 0 d, 6h, 1 d, 3 d, and 5 d post-CFA or saline injection, which was recorded as the average result of 3 trials for each hind paw. The interval time between PWL measurements taken with the ipsilateral and contralateral paws was 5 minutes. So, again PWL measurements of the rats in the DMSO and roscovitine-pretreatment groups were taken by the same method. The behavioral results showed that the heat hyperalgesia induced by CFA significantly decreased from 6h to 5 d and reversed by roscovitine, which is in accordance with our previous studies [21].

### Immunofluorescence

4% paraformaldehyde was injected into the animals which were deeply anesthetized with sodium pentobarbital (> 100 mg/kg. i.p). Tissues collected from ipsilateral segments L4-L6 of the DRG and spinal cord were immediately cryoprotected in 30% sucrose, and sectioned transversely at 16 µm thickness within a cryostat afterwards. The tissue sections were embedded with Tissue-Tek (Sakura Finetek, Torrance, CA, USA) and frozen in dry ice powder. The cross sections were sliced into 16 µm thick sections at -28°C using a cryostat. Using double immunofluorescence for the examination, sections were incubated overnight at 4°C with appropriate antibodies (anti-Cdk5 1:200, ab115812, and anti-VGLUT2, 1:200, ab79157; Abcam, Cambridge, MA, USA), followed by a second overnight incubation at 4°C with the corresponding secondary antibodies (1:200; Invitrogen, Carlsbad, CA, USA). An Axiovert/LSM 510 confocal scanning microscope (Carl Zeiss Microimaging, Inc., Germany) was used to exam the double-stained sections. After that, these sections were washed and the fluorescence-labeled secondary antibodies (Goat anti-Mouse; goat anti-rabbit) were used for the 1 h incubation at room temperature, respectively and 9 slices from 3 different mice in each group. For each slice, Cdk5-positive and VGLUT2-positive [Cdk5(+)/VGLUT2(+)] cells from 4 randomly selected sections (250 μm × 250 μm) within DRG and spinal cord were manually counted [24].

### Protein extraction and western blot analysis

DRG and spinal cord tissues from the L4-L6ipsilateral sides of each treatment groups were collected respectively and immediately stored in ice-chilled lysis buffer (50 mM Tris, pH 7.4, 150 mM NaCL, 1.5 mM MgCL_2_, 10% glycerol, 1% Triton X-100, 5 mM EGTA, 0.5 µg/ml leupetin, 1 mM PMSF, 1 mM Na_3_VO_4_, 10 mM NAF, and a proteinase inhibitor cocktail). Homogenates were centrifuged at 12,000rpm for 15 min at 4°C, and BCA assay kit (Pierce, Rockland, IL, USA) was used to determine the protein concentrations. Each of the samples containing 30 µg of protein were identified by sodium dodecyl sulfate polyacrylamide gel electrophoresis (SDS-PAGE) to determine the estimated molecular mass of the protein and the gel were then transferred onto PVDF membranes. The membranes were then blocked with 1% bovine serum albumin (BSA) in TBST (50 mM Tris-HCL, pH 7.5; 150 mM NaCl, 0.05% Tween 20) for 1 h at room temperature, and incubated overnight at 4°C with the appropriate primary antibody (anti-VGLUT2, 1:500; Abcam, Cambridge, MA, USA, ab79157; anti-p35/p25, anti-p35, 1:300, Cell Signaling, MA, USA, 2680). Membranes were washed in TBST between each of the incubations. Blots were then incubated with a 1:1000 dilution of horseradish peroxidase (HRP)-conjugated secondary antibody (goat anti-rabbit or -mouse, respectively) for 1.5 h at room temperature. The individual bands were visualized by incubation with enhanced chemiluminescence reagent (Boehringer Mannheim, Indianapolis, IN, USA). All bands were exposure to x-ray film which was later digitally analyzed using NIH Image software, version 1.60.

### Statistical analysis

All data are presented as the mean ± SEM. SAS (version 8.01 software for Windows SAS Institute Inc., Cary, North Carolina, USA) was used to calculate statistical results. The time courses of effects were analyzed by two-way analysis of variance, followed by the Dunnett’s test. VGLUT2 expression was analyzed using one-way analysis of variance, followed by the Student Newman–Keuls’ test and P-value < 0.05 was used to indicate statistical significance.

## Results

### Heat hyperalgesia induced by intraplantar CFA injection was significantly reduced by roscovitine

The effects of roscovitine on heat hyperalgesia induced by injection of CFA into the ipsilateral hind paws of rats was determined by measuring heat hyperalgesia on days 1-5 following CFA injection. Compared to the contralateral hind paw of animals in the same group during the same time span, the average thermal PWL threshold (seconds) of the ipsilateral hind paw was significantly reduced 6 hours and on days 1-5 following CFA injection (**P < 0.01, Figure 1 A, *n* = 6/group). However, no pronounced difference was observed between the ipsilateral and contralateral paws of rat in the control group (P > 0.05, Figure 1B, *n* = 6/group). Furthermore, on days 1 and 3 after CFA injection, the average PWL thresholds of isolateral paws pretreated with roscovitine for 30 min prior to intraplantar CFA injection were remarkably increased, when compared to the ipsilateral paws of animals pretreated with DMSO (**P < 0.01, Figure 1 B, *n* = 6/group).

**Figure 1.**
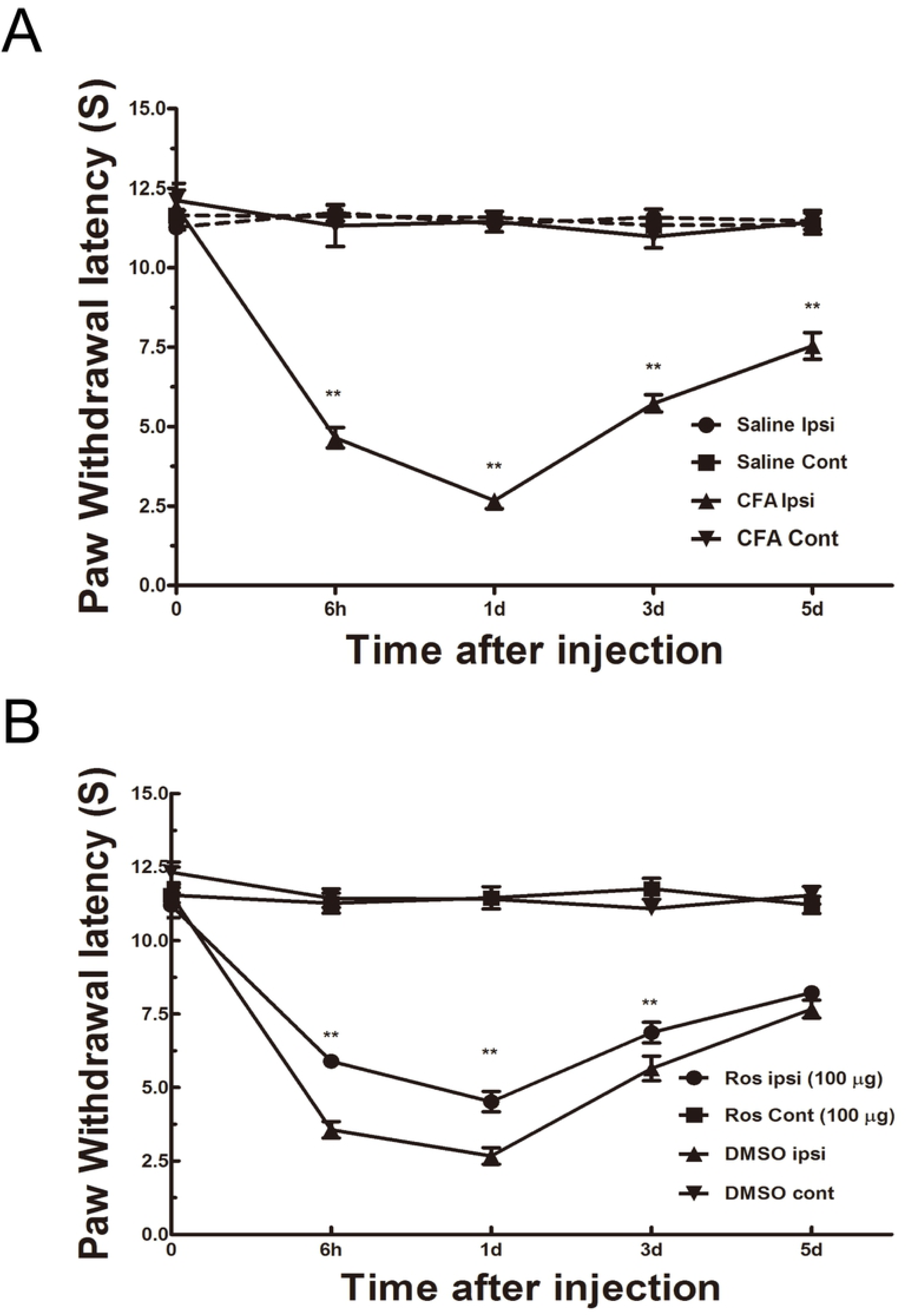
Heat hyperalgesia induced by peripheral injection of CFA was significantly inhibited by intrathecal administration of roscovitine. PWL in response to thermal stimuli significantly decreased following intraplantar injection of CFA. Cont (contralateral) hind paw was injected with saline. Ipsi (ipsilateral) hind paw was injected with CFA. **P < 0.01, n = 6/group (A). Compared to PWLs in the control group pretreated with DMSO, PWL response to thermal stimuli significantly was increased following intrathecal injection of roscovitine 0.5 h prior to CFA injection. Data shown represent the Ave ± SEM, **P < 0.01, n = 6/group (B).

### Upregulated expression of Cdk5/VGLUT2 in DRG neurons

To investigate the possible morphologic relationships between Cdk5 and VGLUT2, light microscopy examinations of the DRG from the L4-L6 segments of spinal cord was to illustrate a dense plexus of immunoreactive elements in the small and medium diameter neuron cells. Compared with the control group injected with saline, coexpression of Cdk5 and VGLUT2 was clearly elevated on both day 1 post-CFA injection (Figure 2 **P < 0.01 *n* = 3/group). These results demonstrated that Cdk5 mediated inflammatory pain by closely interacting with VGLUT2.

**Figure 2.**
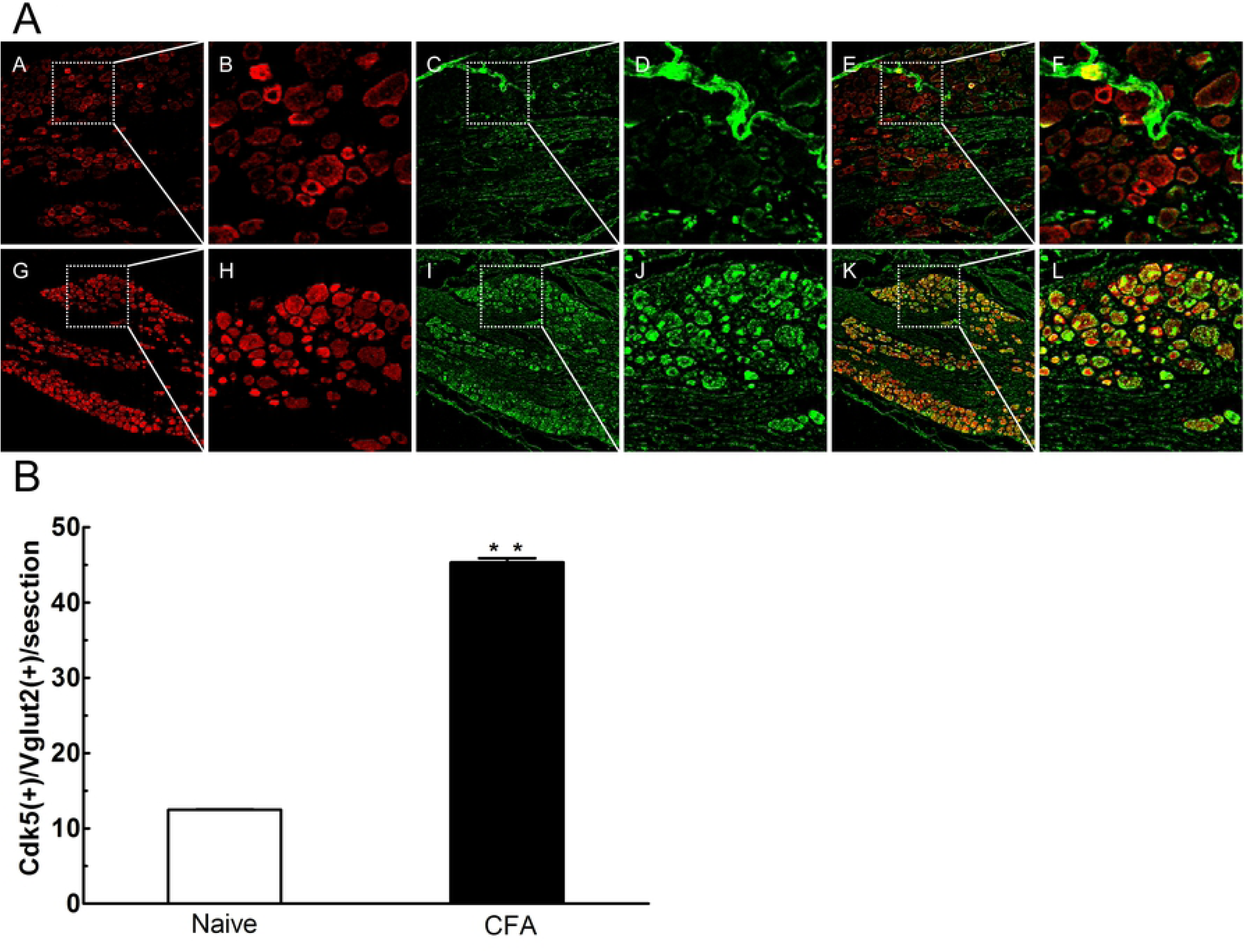
Co-expression of Cdk5 and VGLUT2 was increased in DRG neuronal cells. Double-immunofluorescence staining for Cdk5 (red), VGLUT2(green). Compared with control group (A-F) and day 1(G-L) following intraplantar injection of CFA, co-expression of Cdk5 and VGLUT2 was significantly increased in DRG neurons on day 1 following CFA injection, Data shown represent the Ave ± SEM,*P < 0.05, **P < 0.01, n = 6/group.

### Upregulated expression of Cdk5/VGLUT2 in spinal cord neurons

Subsequently, we observed the co-expression of Cdk5 and VGLUT2 from the segments of L4-L6 spinal cord using double-labeled immunofluorescence. The co-expression of Cdk5/VGLUT2 was preferentially distributed in the gray matter area of spinal cord, which mainly transmitted the pain message from peripheral to central. Compared with the control group challenged with saline, the co-expression of Cdk5 and VGLUT2 was significantly increased on both day 1 post-CFA injection (Figure 3 **P < 0.01 *n* = 3/group).

**Figure 3.**
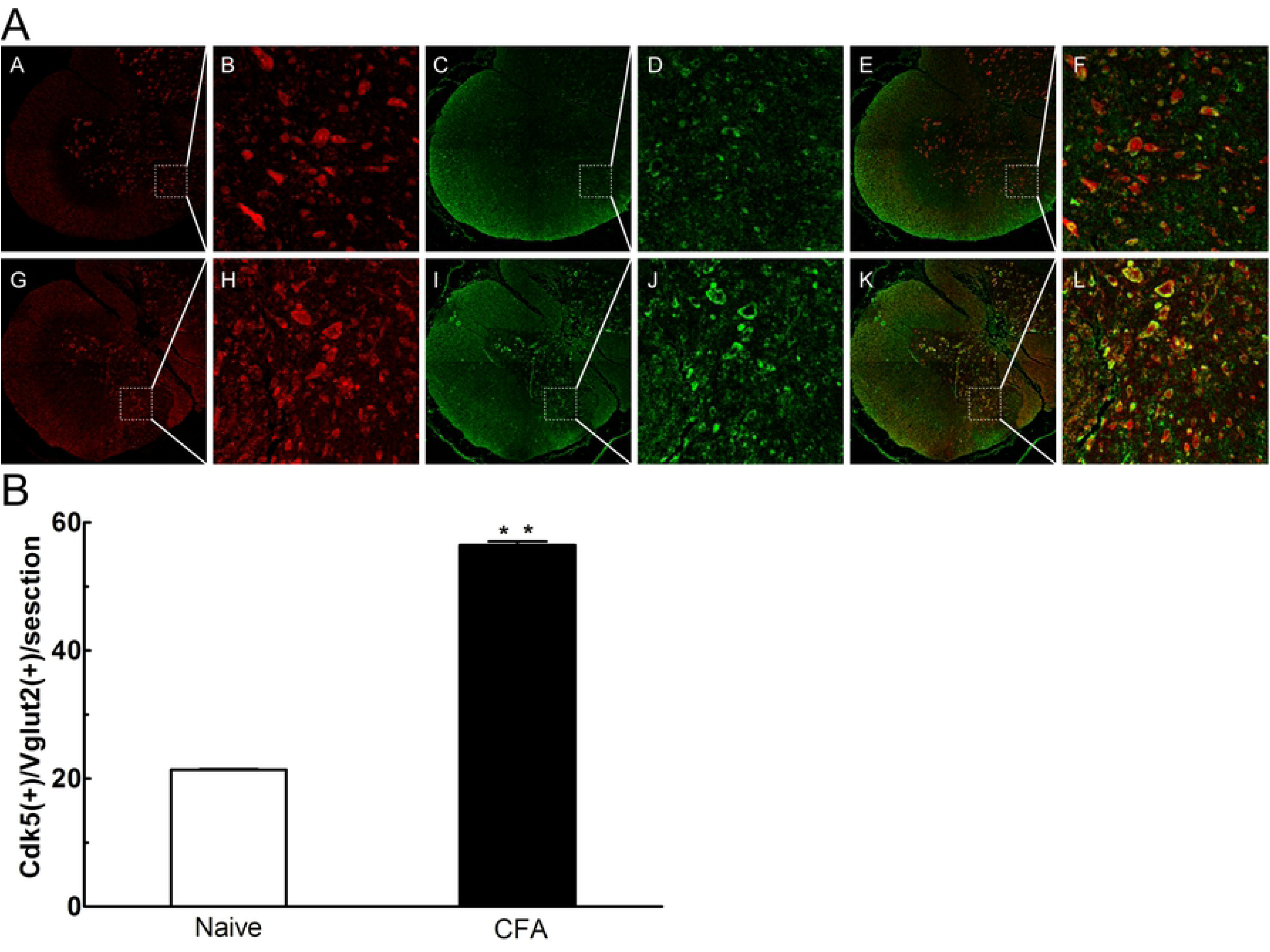
Co-expression of Cdk5 and VGLUT2 was increased in spinal cord neuronal cells. Double-immunofluorescence staining for Cdk5 (red), VGLUT2(green). Compared with control group (A-F) and day 1 (G-L)following intraplantar injection of CFA, co-expression of Cdk5 and VGLUT2 was significantly increased in DRG neurons on day 1 following CFA injection, Data shown represent the Ave ± SEM,*P < 0.05, **P < 0.01, n = 6/group.

### Increased VGLUT2 protein expression in DRG was significantly reduced by roscovitine

We next used western blot analysis to investigateVGLUT2 protein expression in DRG neurons from the L4-L6 ipsilateral sides of spinal cord on days 0, 1, 3, and 5 post-CFA challenges. The results showed that levels of VGLUT2 protein in DRG neurons were significantly increased between days 1 and 3 post-CFA injection (Figure 3 and Figure 5, *P < 0.05, *n* = 4/group). Furthermore, the elevated levels of VGLUT2 protein found between days 1 and 3 were obviously reduced by roscovitine. (Figure 4, **P < 0.01 *n* = 4/group).

**Figure 4.**
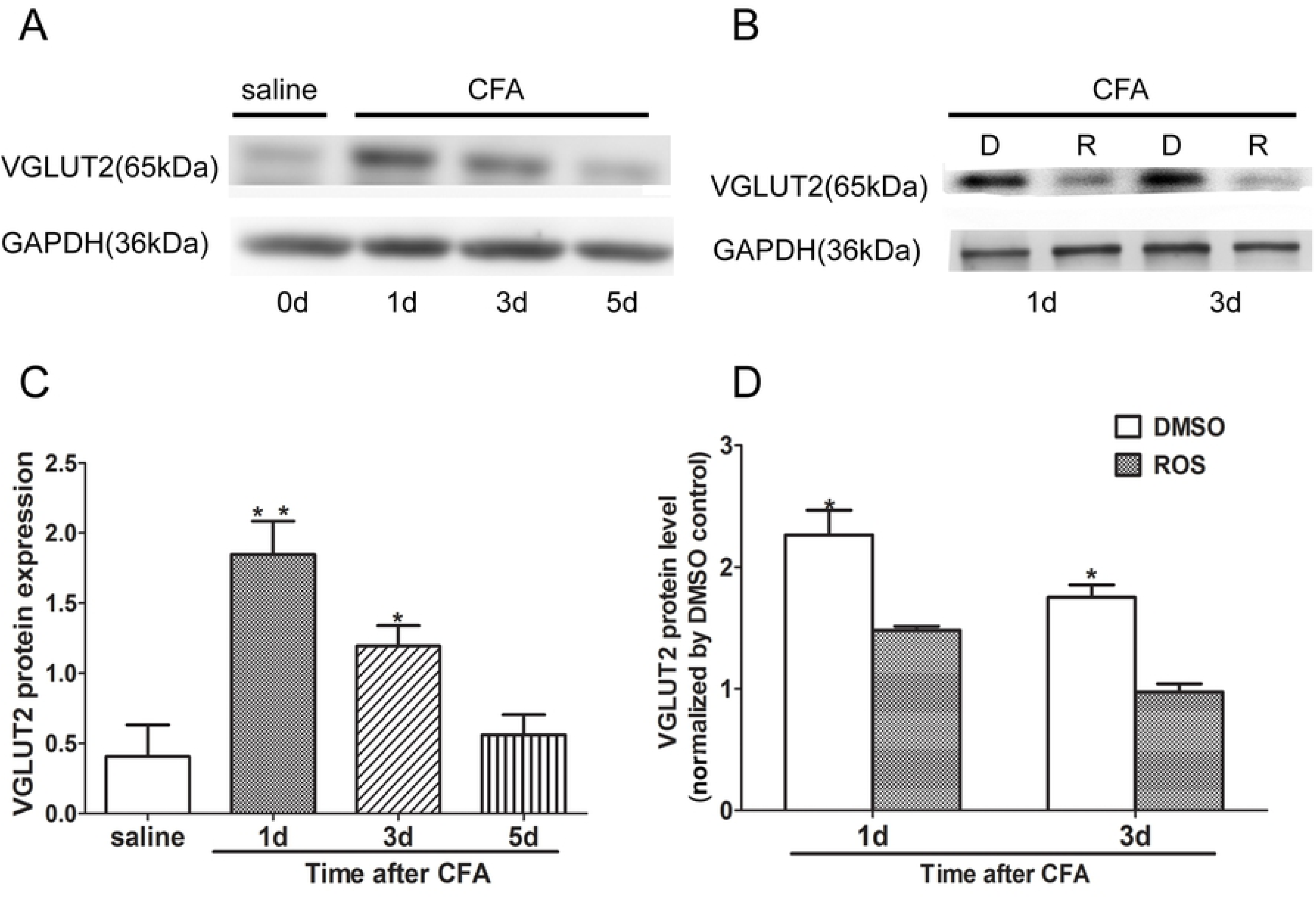
VGLUT2 protein expression was markedly increased in the DRG, and reduced by roscovitine. Compared to the control group, VGLUT2 expression was significantly increased from day 1 to day 3 following intraplantar injection of CFA. *P < 0.05, ** P < 0.01 *n* = 3/group (A). Compared to DMSO-treated controls, the increased expression of VGLUT2 protein was significantly reduced by intrathecal injection of roscovitine between day 1 and day 3. Data shown represent the Ave ± SEM, \**P* < 0.05 *n* = 3/group (B).

**Figure 5.**
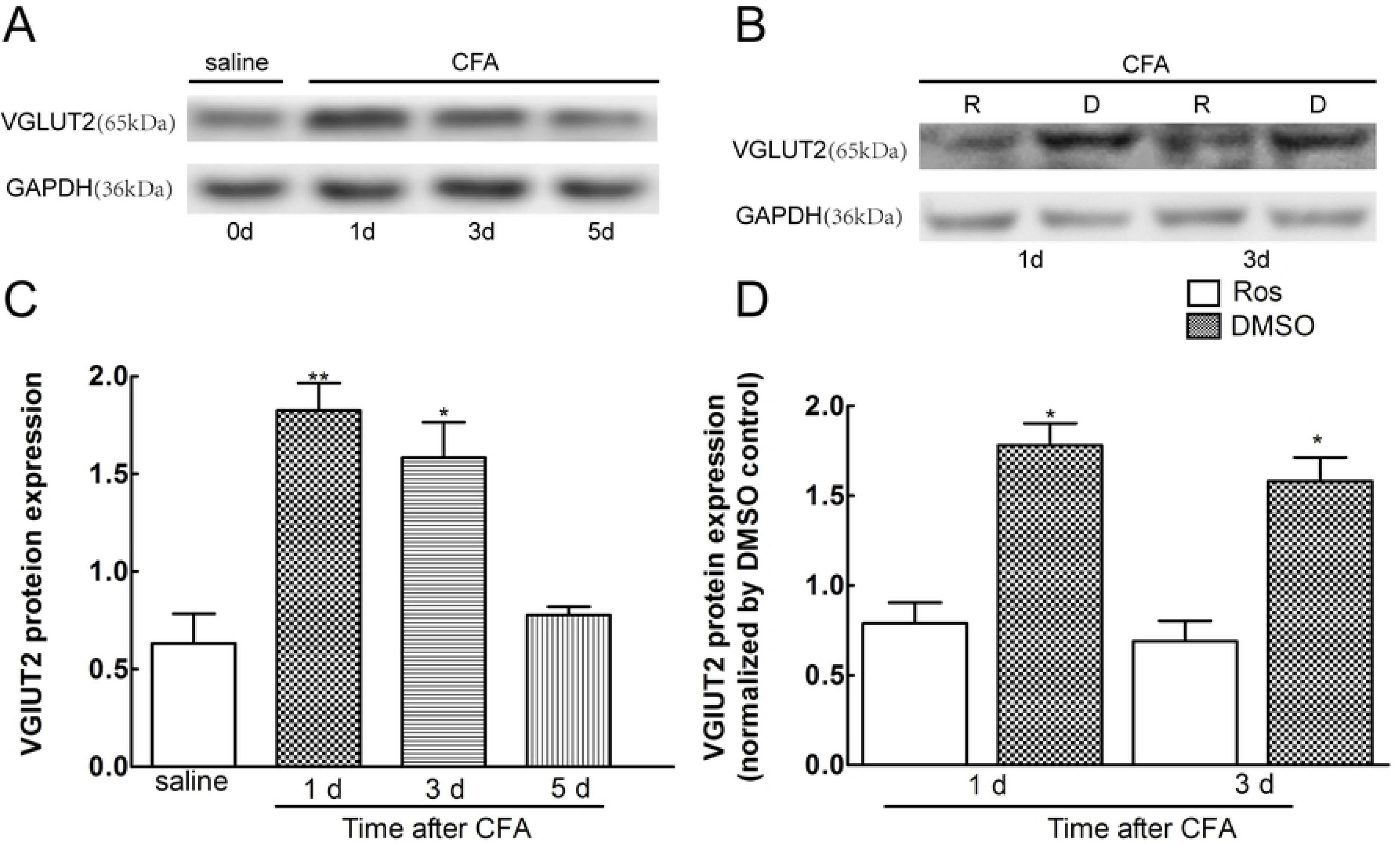
VGLUT2 protein expression was markedly increased in the spianl cord, and reduced by roscovitine. Compared to the control group, VGLUT2 expression was significantly increased from day 1 to day 3 following intraplantar injection of CFA. *P < 0.05, ** P < 0.01 *n* = 3/group (A). Compared to DMSO-treated controls, the increased expression of VGLUT2 protein was significantly reduced by intrathecal injection of roscovitine between day 1 and day 3. Data shown represent the Ave ± SEM, \**P* < 0.05 *n* = 3/group (B).

### Increased VGLUT2 protein expression in spinal cord neurons was significantly reduced by roscovitine

In accodance with the results of VGLUT2 protein expression in DRG, the VGLUT2 protein from the L4-L6 ipsilateral spinal cord neurons challenged with CFA was significantly increased on days 1 and 3 as comapred with the control group. Moreover, the increased VGLUT2 protein was obviously reduced by roscovitine. (Figure 5 **P < 0.01, *n* = 4/group).

### p25 but p35 protein expression in spinal cord neurons was significantly increased by CFA

To varify the role of p25 or p35 as a activator of Cdk5 in mediating the heat hyperalgesia by CFA, the *p25and p35* protein from the L4-L6 ipsilateral spinal cord neurons were exiamed, respectively. p25 protein from the L4-L6 ipsilateral spinal cord neurons challenged with CFA was significantly increased on days 1 and 3 as comapred with the control group. However, p35 remians markedly unchanged as compared with the control group. (Figure 6 **P < 0.01, *n* = 4/group)

**Figure 6.**
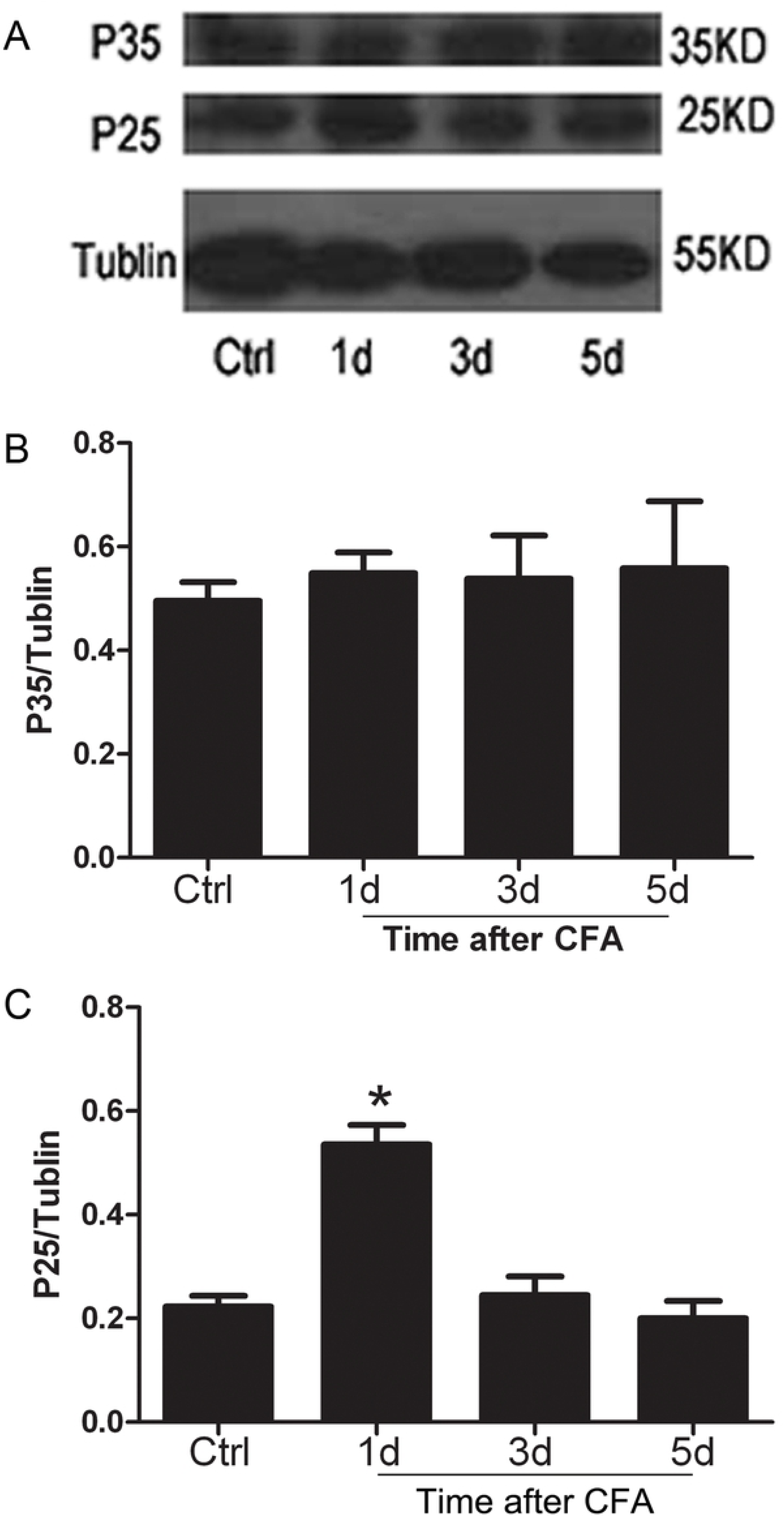
p25 but not p35 was markedly increased in the spianl cord Compared with the control group, p25 expression was significantly increased from day 1 following intraplantar injection of CFA. However, p35 expression was not significantly increased day 1 as compared with the control group. \**P* < 0.05 *n* = 3/group.

### Heat hyperalgesia was significantly inhibited by roscovitine via p25 rather than p35

To explore the pathway inhibiting heat hyperalgesia by roscovitine, we *respectively* detected the protein changing of *p25and p35 after spinal intrathecal adminstration of roscovitine in the inflammation model induced by CFA. Basicaly, in agreement with the results of FIG 6, the expression of p25 was significantly reduced by roscovitine but not p35, suggesting that inhibition of heat hyperalgesia by Cdk5 inhibitor* roscovitine was via inhibiting the activity of p25 not p35. (Figure 7 **P < 0.01, *n* = 4/group)

**Figure 7.**
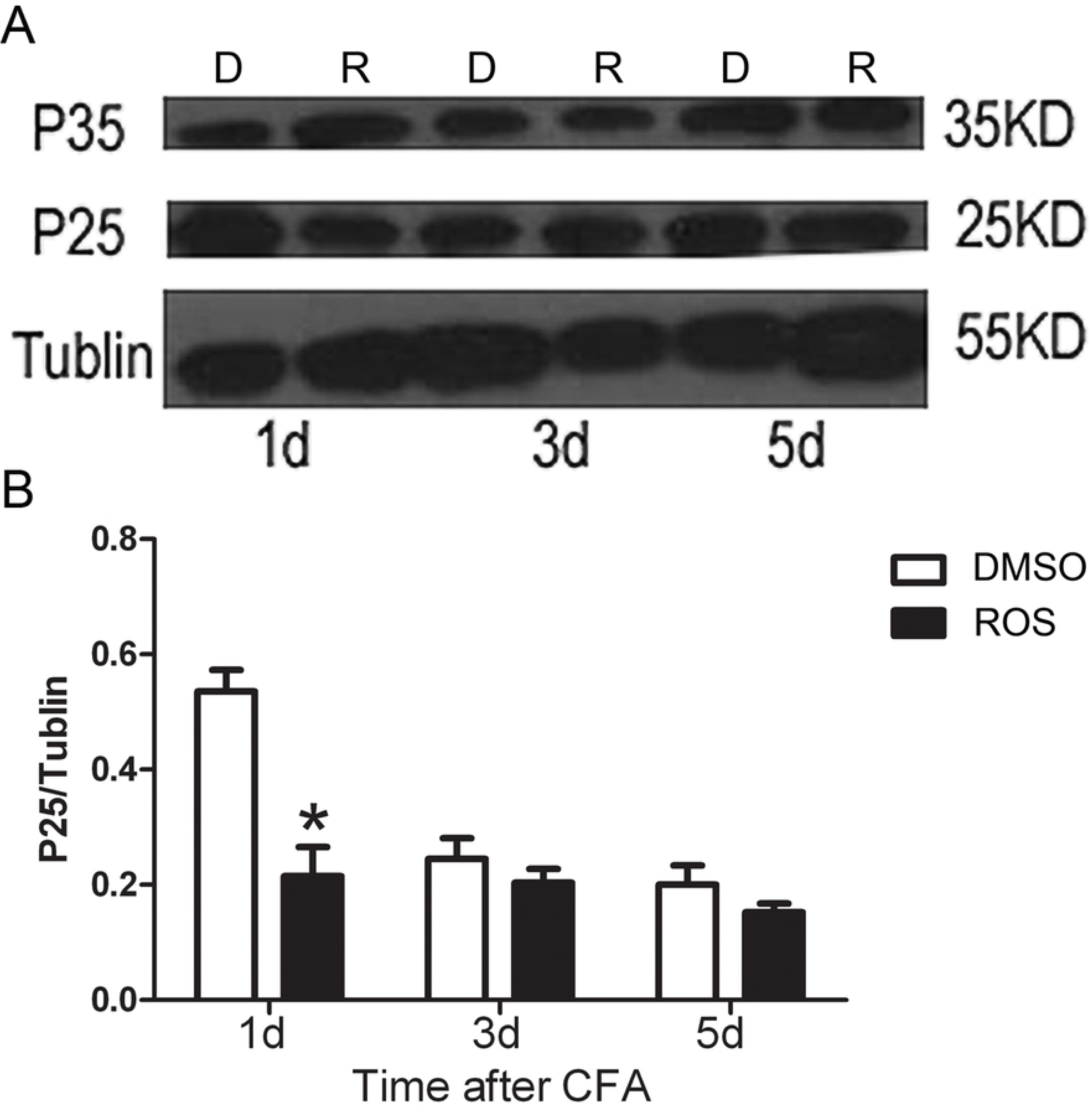
Heat hyperalgesia was significantly inhibited by roscovitine via p25 rather than p35 Compared to DMSO-treated controls, the increased expression of p25 protein was significantly reduced by intrathecal injection of roscovitine between day 1. However, the effects of administration of on p35 protein expression. Data shown represent the Ave ± SEM,\**P* < 0.05 *n* = 3/group (B).

## Discussion

The present study preliminary illustrated that the VGLUT2 signaling pathway was involved in CFA-induced heat hyperalgesia mediated by Cdk5 in the DRG and spinal cord neurons. It is well established that glutamate is the major excitatory neurotransmitter in the CNS, and plays a key role in nociceptive processing in inflammatory and neuropathological pain [1]. Storage of glutamate within vesicles is controlled by VGLUTs, and small DRG neuron cells accumulate 6-fold more glutamine, a precursor in the synthesis of L-glutamate, than larger DRG neurons [25]. As currently determined, the family of VGLUTs consisting of VGLUTs 1-3, is not overlapped, and only VGLUT 2 plays a key role in mediating the heat hyperalgesia induced by inflammation [6, 7]. Previous studies conducted using a CFA-induced inflammatory pain model has shown that the thresholds of heat hyperalgesia in mice with knocked out VGLUT2 were markedly increased compared with wild mice. However, the mechanisms by which VGLUT2 mediates inflammation-induced heat hyperalgesia remains to be explored in every detail.

Considerable evidence demonstrated that Cdk5 acts a key role in mediating heat hyperalgesia induced by inflammation [13-16]. Moreover, previous results obtained by other investigators suggest that Cdk5 is a key regulator in mediation of neurotransmitters, including glutamate in the CNS [17]. Our previous studies demonstrated that synaptophysin, an important presynaptic vesicle membrane protein which mediates release of neurotransmitters, was involved in heat hyperalgesia mediated by Cdk5, suggesting that Cdk5 may mediate heat hyperalgesia induced by inflammation by controlling the release of neurotransmitters. A recent study further indicated that VGLUT2 was invovled in the trafficking of synaptic vesicles by interacting with synaptophysin at synaptic boutons [19], which further show that VGLUT2 mediates the heat hyperalgesia via Cdk5 pathway. In addition, another study made it clear that the expression of VGLUT1 and VGLUT2 gene was strongly reduced in synapsin II (another key vesicle protein involved in release of neurotransmitters) knockout mice [18]. Thus, we examined the possible linkage between Cdk5 and VGLUT2 in mediating heat hyperalgesia induced by CFA using our current model.

In our research, the immunohistochemistry results showed that the co-expression of Cdk5 and VGLUT2 was obviously in small and medium-sized neuronal cells of the DRG and spinal cord. It had been established that small- and medium-sized neuronal cells of the DRG and spinal cord are important mediators of inflammation and neuropathologic pain. Moreover, the small and medium-sized neuronal cells of the DRG and spinal cord have been shown to express a variety of nociceptive ion channels and receptors. The results from our current study align with findings from previous studies, in that Cdk5 and VLGUT2 are prominently distributed in the small and medium-sized neuronal cells of the DRG and spinal cord. More importantly, our results demonstrate that Cdk5 and VLGUT2 are co-expressed in small and medium-sized neuronal cells of the DRG and spinal cord. Additionally, the co-expression of Cdk5 and VGLUT2 was found to be significantly increased between days 1 and 3 following plantar injection of CFA, as compared with the control group of rats that was challenged with saline. The increased co-expression of Cdk5 and VGLUT2 lasted for more than 3 days, until day 5, which suggests a key role of these molecular factors in the induction stage of chronic inflammatory pain. Subsequently, our study explored the effects of Cdk5 on VGLUT2 by administering roscovitine via intrathecal injection on day 1 and day 5 prior to CFA injection, which allowed us to observe the changes in protein of VGLUT2 under the heat hyperalgesia condition.

Previous studies conducted using a CFA-induced inflammatory pain model demonstrated that thresholds of heat hyperalgesia in mice with VGLUT2 knockout in the DRG were significantly increased compared to thresholds in wild-type mice, suggesting that VGLUT2 plays a key role in mediating the heat hyperalgesia induced by peripheral injection of CFA. In our study, both VGLUT2 and Cdk5 expression in the DRG and spinal cord were significantly increased between days 1 and 3 following CFA injection, and returned to baseline levels on day 5. In the current study, an intrathecal injection of roscovitine was used to determine whether endogenous VGLUT2 was inhibited by roscovitine in DRG and spinal cord neurons. The results showed that increased VGLUT2 protein expressions were significantly reduced by roscovitine. Although the activities of Cdc2, Cdk2, and Cdk5 kinase are all inhibited by roscovitine [26], our previous findings suggested that in adult rats, roscovitine mainly affects the activity of Cdk5, and does not affect the activity of other Cdks [21]. In addtion, p25 but not p35 as the activator of Cdk5 contributed to the CFA-induced heat hyperalgesia. It was well established that Cdk5’ activity mainly depends on its activator. Thus, our data suggests that inhibition of Cdk5 kinase activity by p25 led to the decreases in VGLUT2 protein expression observed in our current model. Taken together, our findings suggest that in our current model, Cdk5 functions as the key regulator of VGLUT2.

It is important to note that the results from our model system failed to pinpoint exactly how Cdk5 mediates the neurotransmitters release, e.g. glutamate (excitatory) interaction with VGLUT2, or how Cdk5 mediates the synaptic vesicles transportation, e.g. by closely interacting with VGLUT2. In addtion, we only used VGLUT2 antiboy for immunohistochemistry and protein expression changing in our model to show the role of pain transmission mediated by VGLUT2, lacking the more specific method in our model. Future experiments will be designed to determine the mutual relationship between Cdk5 and VGLUT2 in mediating the release of excitatory neurotransmitters of glutamate in this model of knockout of VGLUT2 as contorl.

Our studies demonstrate that Cdk5 mediates CFA-induced heat hyperalgesia by regulating VGLUT2 protein expression in DRG and spinal cord neurons. Increased co-expression of Cdk5 and VGLUT2 was observed in small and medium-sized neuronal cells of the DRG and spinal cord induced by CFA. Spinal administration of roscovitine markedly alleviated heat hyperalgesia induced by CFA. Furthermore, increased VGLUT2 protein expression in DRG and spinal cord neurons was reduced by roscovitine, suggesting that Cdk5/p25, as the upstream regulator of VGLUT2, plays an important role in inducing and developing the early stage of chronic inflammatory pain. Consequently, we propose that severing the linkage between Cdk5/p25 and the VGLUT2 signaling pathway may present a promising therapeutic strategy for diminishing inflammatory pain.

## Acknowledgements

The work supported by the Foundation of Technology Bureau of Hang Zhou (grant no. 20150733Q01), the Technology Plan of Chinese Traditional Medicine of Zhejiang Province (grant no. 2014ZB067 and 2015ZA136), the Zhejiang Provincial Natural Science Foundation of China (grant no. Y14H090005), the National Natural Science Foundation, Beijing, P.R. China(grant no. 81771403), the Program of New Century 131 outstanding young talent plan top-level of Hang Zhou and the Program of Medical and Health Technology Projects of Zhejiang Province (grant no. 2017RC012) to Dr. Hong-Hai Zhang, the National Natural Science Foundation, Beijing, P.R. China (grant no. 81401072) to Dr. Gonglu Liu, the National Natural Science Foundation, Beijing, P.R. China (grant no. 81401072) to Dr. Zhiyou Peng(grants No. 81603198), the Health Science and Technology Plan of HangZhou (grant no. 2017A07) to Dr.Xian-Zhe-Luo, the Health Science and Technology Plan of HangZhou and the Health Science and Technology Plan of Zhejiang Province (grant no. 2018A17 and 2019ZD015) to Dr. ShouJun-Tao and the Health Science and Technology Plan of Zhejiang Province (grant no. 2018KY570) to Dr.Qing-Hua Li. We thank for the supporting of Shanghai Sunbio Bio-Medicine Technology Co.,Ltd.

## Author Contributions statement

Conceived and designed the experiments: HongHai Zhang, ZhiYing Feng

Performed the experiments: YuWen Tang, ZhiYou Peng, ShouJun-Tao, Jianliang Sun, WenYuan Wang, XueJiao Guo

Analyzed the data: Peng Xu, Qing Hua Li

Contributed reagents/materials/analysis tools: GongLu Liu, XianZhe-Luo, Yuan Chen, Yue Shen, HaiXiang Ma

